# Optimising Renewal Models for Real-Time Epidemic Prediction and Estimation

**DOI:** 10.1101/835181

**Authors:** KV Parag, CA Donnelly

## Abstract

The effective reproduction number, *R*_*t*_, is an important prognostic for infectious disease epidemics. Significant changes in *R*_*t*_ can forewarn about new transmissions or predict the efficacy of interventions. The renewal model infers *R*_*t*_ from incidence data and has been applied to Ebola virus disease and pandemic influenza outbreaks, among others. This model estimates *R*_*t*_ using a sliding window of length *k*. While this facilitates real-time detection of statistically significant *R*_*t*_ fluctuations, inference is highly *k* -sensitive. Models with too large or small *k* might ignore meaningful changes or over-interpret noise-induced ones. No principled *k* -selection scheme exists. We develop a practical yet rigorous scheme using the accumulated prediction error (APE) metric from information theory. We derive exact incidence prediction distributions and integrate these within an APE framework to identify the *k* best supported by available data. We find that this *k* optimises short-term prediction accuracy and expose how common, heuristic *k* -choices, which seem sensible, could be misleading.

## Introduction

The time-series of infected cases (infecteds) observed over the course of an infectious disease epidemic is known as an incidence curve or epi-curve [1]. At the population level, these curves offer prospective insight into the spread of a disease by informing on the effective reproduction number, which defines the average number of secondary infections induced by a primary one [2]. This reproduction number, denoted *R*_*t*_ at time *t*, is an important prognostic of the behaviour of an epidemic. If *R*_*t*_ > 1, for example, we can expect and hence prepare for an exponentially increasing incidence, whereas if *R*_*t*_ < 1, we can be reasonably confident that the epidemic has been arrested [2]. Reliably estimating meaningful changes in *R*_*t*_ is an important problem in epidemiology, since it can forewarn about the growth rate of an outbreak and signify the level of control effort that must be initiated or sustained [3] [4].

The renewal model [5] is a popular approach for inferring salient fluctuations in *R*_*t*_ and has been used to predict Ebola virus disease case counts and assess the transmission potential of pandemic influenza and Zika virus, among others [6] [7] [3]. The renewal model may be applied retrospectively to understand the past behaviour of an epidemic or prospectively to gain real-time insight into ongoing outbreak dynamics [8] [5]. We only consider the latter here. Model selection metrics for the former application were investigated in [9]. The prospective approach approximates *R*_*t*_ with a piecewise-constant function i.e. *R*_*t*_ is considered constant or stable over some sliding window of *k* time units (e.g. days or weeks) into the past, beyond which a discontinuous change is assumed [8].

This formulation models the non-stationary nature of epidemics, capturing the idea that different dynamics are expected during distinct phases (e.g. onset, growth, control) of the epidemic lifetime. The window length, *k*, is effectively a hypothesis about the stability of *R*_*t*_ underpinning the observed epi-curve and is critical to reliably characterising the epidemic because it controls the time-scale over which *R*_*t*_ fluctuations are deemed significant. Too large or small a *k*-value can respectively lead to over-smoothing (which ignores important changes) or to random noise being misinterpreted as meaningful. Inferring under the wrong *k* can appreciably affect our understanding of an epidemic, as observed in [5], where case reproduction numbers (a common *k*-grouping of *R*_*t*_) were found to over-smooth significant changes in HIV transmission, for example.

Surprisingly, no principled method for optimally selecting *k* exists. Current best practice either relies on heuristic choices (including visual trial and error) [6] or provides implicit, minimum thresholds for *k* [8]. Here we adapt metrics from information theory to develop a simple yet rigorous framework for optimising *k*. Specifically, we analytically derive the posterior predictive incidence distribution of the renewal model and then minimise its accumulated prediction error (APE) [10] over the space of possible *k*, to obtain *k*\*. The APE is a formal, data-driven, model selection strategy that identifies the *k*\* most justified by the available incidence data by optimising for short-term predictive accuracy [11]. The renewal model with this *k*\* best explains the observed epi-curve.

The APE metric is valid for all sample sizes, is easily computed for models of arbitrary dimension and is strongly linked to Bayesian model selection (BMS), cross validation and minimum description length (MDL) [12] [13]. Unlike many standard model selection approaches, the APE accounts for parametric complexity, which describes how functional dependencies among parameters matter [14]. Ignoring parametric complexity is known to promote suboptimal renewal model selection performance [9]. The APE metric deems the model that best predicts unseen data from the same underlying process, and not the one that best fits existing data, as optimal and of justified complexity [15].

Our APE-based approach should therefore pinpoint and characterise only those reproduction number fluctuations that are key to achieving reliable short-term incidence predictions. The performance, speed and computational ease of our approach makes it suitable for real-time forecasting and model selection. It could serve as a stand-alone computational tool or be integrated within existing real-time frameworks, such as in [4], to provide emerging insights into infectious transmission dynamics, or to assess the prospective efficacy of implemented interventions (e.g. vaccination or quarantine). Public health policy decisions or preparedness plans based on improperly specified *k*-windows could be misinformed or overconfident. Our approach will hopefully limit these risks.

## Results

### Problem Definition

Let the incidence or number of infected cases in an epidemic at present time *t*, be *I*_*t*_. The incidence curve is a historical record of these case counts from the start of the outbreak [1], and is summarised as 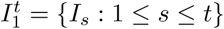, with *s* as a time-indexing variable. For convenience, we assume that incidence is available on a daily scale so that 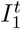 is a vector of *t* daily counts (though in general weeks or months could be used instead). The associated effective reproduction number and total infectiousness of the epidemic are denoted *R*_*t*_ and Λ_*t*_, respectively.

Here *R*_*t*_ is the number of secondary cases that are induced, on average, by a single primary case at *t* [2], while Λ_*t*_ measures the cumulative impact of past cases, from the epidemic origin at *s* = 1, on the present. The generation time distribution of the epidemic, which describes the elapsed time between a primary and secondary case, controls Λ_*t*_ (see Methods). In the problems we consider, 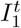 and 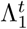 form our data, the generation time distribution is assumed known, and 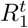 or some grouping of this vector are the parameters to be inferred.

The renewal model [5] [6] derives from the classic renewal epidemic transmission equation [26], and defines the Poisson distributed (Poiss) relationship *I*_*t*_ ∼ Poiss (*R*_*t*_Λ_*t*_). This means that 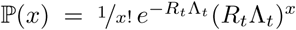, with *x* indexing possible *I*_*t*_ values. This formulation assumes maximum epidemic nonstationarity (i.e. that transmission statistics change every time unit) and hence only uses the most recent data (*I*_*t*_, Λ_*t*_) to infer the current *R*_*t*_. While this maximises the fitting flexibility of the renewal model, it often results in noisy and unreliable estimates that possess many spurious reproduction number changes [6]. Consequently, grouping is employed. This hypothesises that the reproduction number is constant over a *k*-day window into the past and leads to a piecewise-constant function that separates salient fluctuations (change-points) from negligible ones (the constant segments) [8] [9].

We use *R*_*τ*(*t*)_ to indicate that the present reproduction number to be inferred is stationary (constant) over the last *k* points of the incidence curve i.e. the time window *τ* (*t*) := {*t, t* − 1, …, *t* − *k* + 1}. The data used to estimate *R*_*τ*(*t*)_ is then 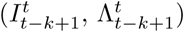. This construction allows us to filter noise and increase estimate reliability but elevates bias. In the Methods we show precisely how grouping achieves this bias-variance trade-off. Choosing *k* therefore amounts to selecting a belief about the scale over which the epidemic statistics are meaning-fully varying and can significantly influence our understanding of the dynamics of the outbreak. Thus, it is necessary to find a principled method for balancing *k*. Ultimately, we want to find a *k*, denoted *k*\*, that (i) is best supported by the epi-curve and (ii) maximises our confidence in making real-time, short-term predictions that can inform prospective epidemic responses.

This is no trivial task as *k*\* would be sensitive to both the specifically observed stochastic incidence of an epidemic and to how past infections propagate forward in time. We solve (i)-(ii) by applying the accumulated prediction error (APE) metric from information theory [10], which values models on their capacity to predict unseen data from the generating process instead of their ability to fit existing data [11]. The properties and mathematical definition of the APE are provided in the Methods. The APE uses the window of data preceding time 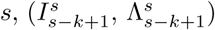, to predict the incidence at *s* + 1 and assigns a log-score to this prediction. This procedure is repeated over *s* ≤ *t* and for possible *k*-values. The *k* achieving the minimum cumulated log-score is deemed optimal. Fig. 1 illustrates the APE for selecting between two windows *k*_1_ and *k*_2_.

**Figure 1:**
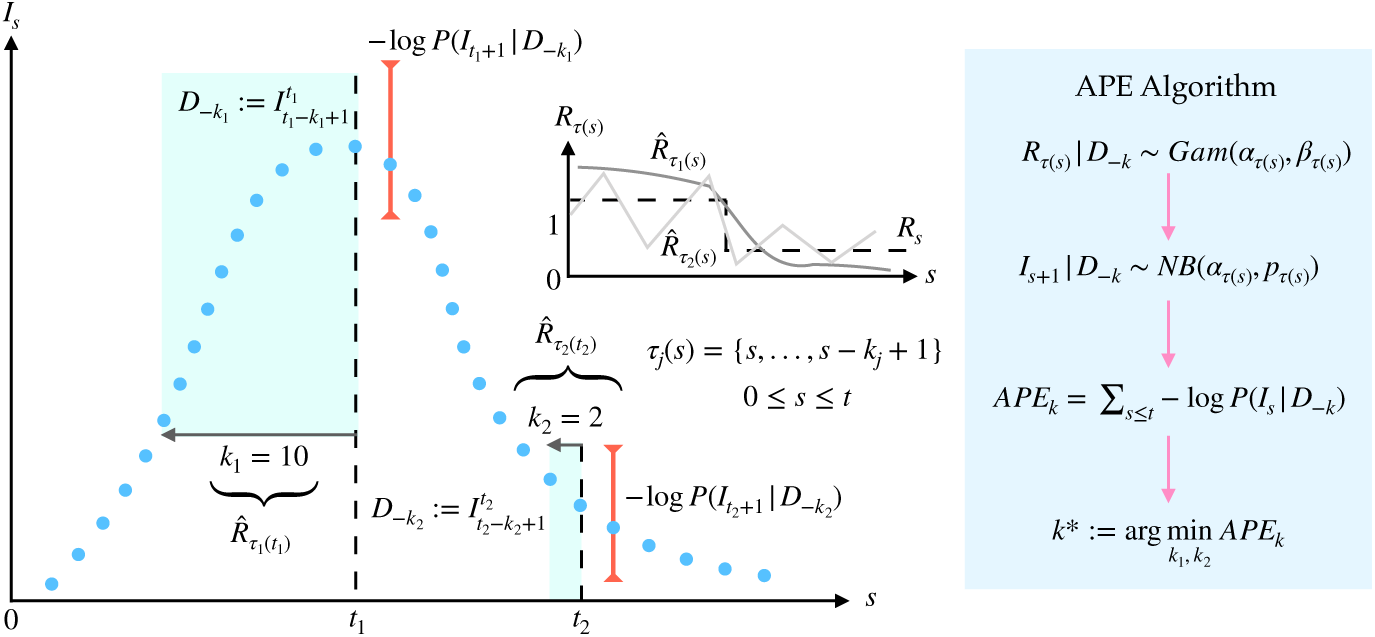
Window length selection using APE. An observed incidence curve (blue dots) is sequentially and causally predicted over time *s* ≤ *t*, using effective reproduction number estimates based on two possible windows lengths of *k*_1_ and *k*_2_ (green shaded). The true reproduction number (*R*_*s*_) is drawn in dashed black (inset). Its estimates for each window length are 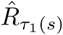 (dark grey) and 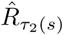 (light grey). Large windows (*k*_1_) smooth over fluctuations. Small ones (*k*_2_) recover these changes but are noisy. The APE assesses *k*_1_ and *k*_2_ via the log-loss of their sequential predictions (red error bars shown for predictions at *t*_1_ and *t*_2_ under *k*_1_ and *k*_2_ respectively). The *k*-window with the smaller APE is better supported by this incidence curve. The algorithm parameters are defined in Results.

### Renewal Model Prediction

To successfully adapt APE, we require the posterior predictive incidence distribution of the renewal model, which at time *s* is 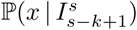 (see Eq. (9)), with *x* indexing the space of possible one-step-ahead predictions at *s* + 1, and *Î*_*s*+1_ = 𝔼[*x*]. We assume a gamma conjugate prior distribution on *R*_*τ*(*s*)_ as in [8]. For some hyperparameters *a* and *c* this is *R*_*τ*(*s*)_ ∼Gam (*a*, 1*/c*), with Gam as a standard shape-scale parametrised gamma distribution. The posterior distribution of *R*_*τ*(*s*)_ is 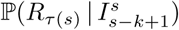 and is described in Eq. (1) [8].

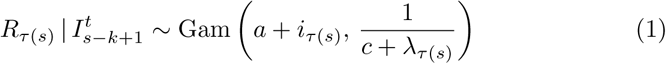

For convenience we define *α*_*τ*(*s*)_ := *a* + *i*_*τ*(*s*)_, *β*_*τ*(*s*)_ := 1*/c*+*λ*_*τ*(*s*)_ with *i*_*τ*(*s*)_ and *λ*_*τ*(*s*)_ are the sum of incidence (*I*_*s*_) and total infectiousness (Λ_*s*_) over the window *τ* (*s*) (see Methods). If a variable *y* ∼ Gam(*α, β*) then 𝕡 (*y*) = *y*^*α*−1^*e*^−*y/β*^/*β*^*α*^ Γ(*α*) and 𝔼 [*y*] = *αβ*. The posterior mean estimate is therefore 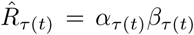. Applying Bayes formula and marginalising yields the posterior predictive distribution of the number of infecteds at *s* + 1 as in Eq. (2).

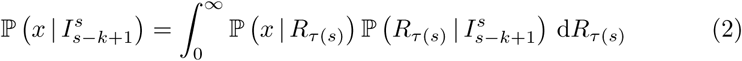

In Eq. (2) we used the conditional independence of future incidence data from the past epi-curve, given the reproduction number to reduce 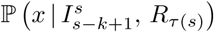 to 𝕡(*x* | *R*_*τ* (*s*)_), which expresses the renewal model relation *x* ∼ Poiss(*R*_*τ*(*s*)_Λ_*s*+1_). As Λ_*s*+1_ only depends on 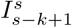 (see Methods) there are no unknowns. Solving using this and Eq. (1) gives 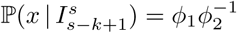 with *ϕ*_1_, *ϕ*_2_ from Eq. (3).

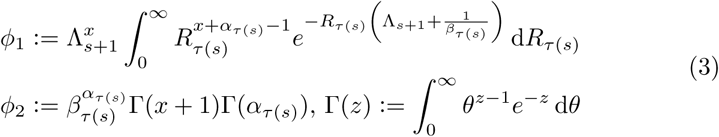

Since ∫ *θ*^*z*−1^*e*^−*yθ*^ d*θ* ≡ Γ(*z*)*/y*^*z*^ the integral in *ϕ*_1_ simplifies to 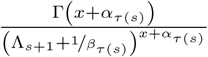. Some algebra then reveals the negative binomial (NB) distribution of Eq. (4).

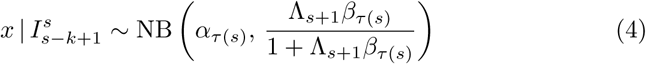

For simplicity we define 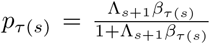 (see Fig. 1). If some variable *y* ∼ NB(*α, p*) then 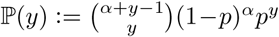 and 𝔼[*y*] = *pα/*1−*p*. Eq. (4) is a key result and relates to a framework developed in [4]. It completely describes the one-step-ahead prediction uncertainty and has mean 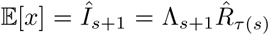, and variance 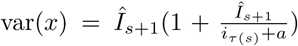 If *i*_*τ*(*s*)_ ≫ *a* and *λ*_*τ*(*s*)_ ≫ *c* then 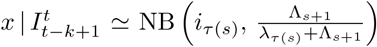. These relations explicate how the current estimate of *R*_*τ*_ (*s*) influences our ability to predict upcoming incidence points. Setting *s* = *t* in all the above expressions will give the prediction statistics for the next (unobserved) time-point beyond the present.

Importantly, Eq. (4) controls the APE metric through the shape of its distribution. We explicitly compute this to derive Eq. (5), with *I*_*s*+1_ as the (true) observed incidence at time *s* + 1, which is evaluated within the context of the predictive space of *x*, and 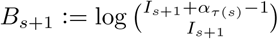 as a binomial term.

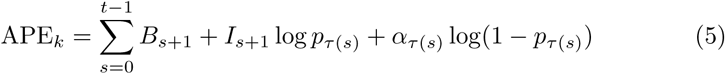

Eq. (5) is easy to evaluate using the in-built NB routines of many software. When computing directly, the most difficult term is *B*_*s*+1_ when *I*_*s*+1_ and *α*_*τ*(*s*)_ are large. In these cases Stirling approximations may be applied. The renewal model APE metric offers a simple means of finding *k*\* := arg min_*k*_ APE_*k*_, the window length that optimises one-step-ahead predictive performance.

### Optimal Window Selection

We apply our APE metric to select *k*\* for various epidemic examples, which examine reproduction number profiles for large outbreaks (Fig. 2a–Fig. 2c), small ones (which are more difficult to estimate: Fig. 2d–Fig. 2e), and a random one (Fig. 2f). We simulated epidemics for *t* = 200 days using a generation time distribution analogous to that used for Ebola virus disease predictions in [16]. We investigated a window search space of 2 ≤ *k* ≤ *t/*2 and computed the APE at each *k*, over the epidemic duration (1 ≤ *s* ≤ *t*). Reproduction number estimates, 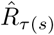, and incidence predictions, 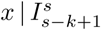, are presented on the left and right panels in Fig. 2. Predictions over the first *s* < *k* times use all *s* data points.

**Figure 2:**
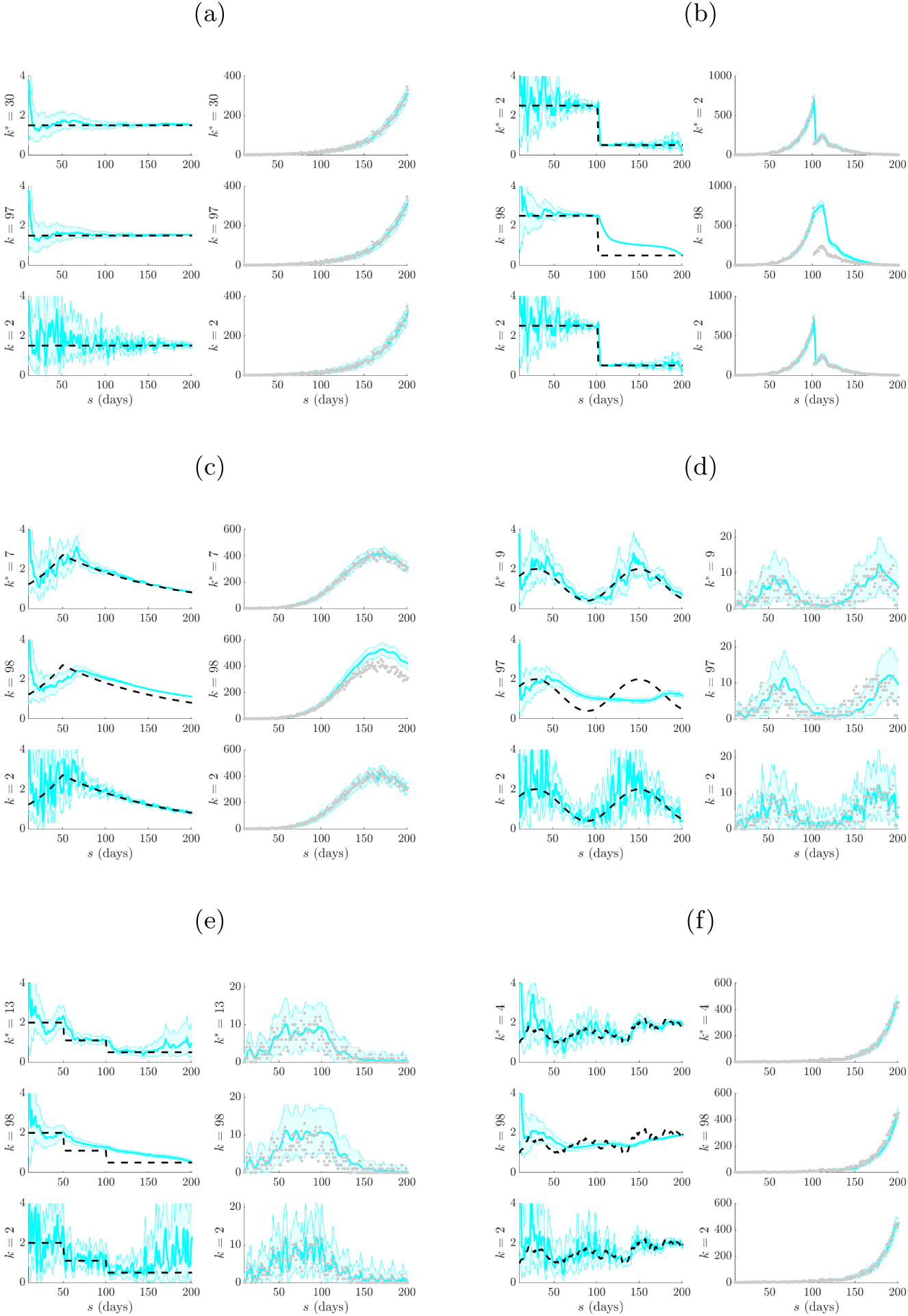
Selection under time-varying reproduction numbers. Left graphs of each panel compare 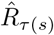 estimates (cyan with 95% confidence intervals) at the APE window length *k*\* to those when *k* is set to the upper or lower limits of its search space. Right graphs give corresponding one-step-ahead predicted incidence curves (cyan with 95% prediction intervals). Dashed lines are the true *R*_*s*_ numbers (left) and dots are the true *I*_*s*_ counts (right). The panels examine (a) stable (constant), (b) steeply changing, (c) exponentially rising and decaying, (d) periodically varying, (e) piecewise falling and (f) random (filtered white noise) changes in *R*_*s*_.

We find that the APE metric balances 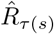 estimate accuracy against one-step-ahead predictive coverage (i.e. the proportion of time that the true *I*_*s*+1_ lies within the 95% prediction intervals of 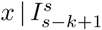) for a range of *R*_*s*_ dynamics (top graphs of Fig. 2). This dual optimisation is of central interest to this swork. Moreover, vastly different *I*_*s*+1_ predictions and 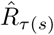 estimates can result when *k*-misspecification occurs (middle and bottom graphs in Fig. 2). This can be especially misleading when attempting to identify the significant changes in epidemic transmissibility and can support substantially different beliefs about the progression of a pathogen. Estimation and prediction performance also depend on the actual incidence at any time, with small *I*_*s*_ values necessitating wider confidence intervals in 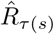 at any *k* (see Eq. (8) of Methods).

When *k* is unjustifiably large not only do we observe systematic prediction and estimation errors, but alarmingly, we tend to be overconfident in them. When *k* is too small, we infer rapidly and randomly fluctuating 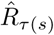 values, which sometimes can deceptively underlie reasonably looking incidence predictions. Consequently, optimal *k*-selection is integral for trustworthy inference and prediction. Observe that small *k*, which implies a more complex renewal model (i.e. there are more parameters to be inferred), does not generally result in better causally predictive one-step-ahead incidence predictions. Had we instead naively picked the *k* that best fits the existing epi-curve, then the smallest *k* would always be favoured (overfitting) [9].

We emphasize and validate the predictive performance of APE in Fig. 3. This shows, for all simulated examples from Fig. 2, that minimising the APE also approximately minimises the percentage of the time *s* ≤ *t* that true incidence values fall outside the 95% prediction intervals of Eq. (4). This validates the APE approach and evidences its proficiency at optimising for short-term forecasting accuracy in real time. We recommend using APE to define *k*\* for an existing epi-curve up to the present *t*. Applying Eq. (4) with this *k*\* should then best predict the number of cases on the (*t* +1)^th^ day. This entire procedure should be repeated for later forecasts, with *k*\* being progressively updated. The behaviour of *k*\* as data accumulate is investigated in the next section.

**Figure 3:**
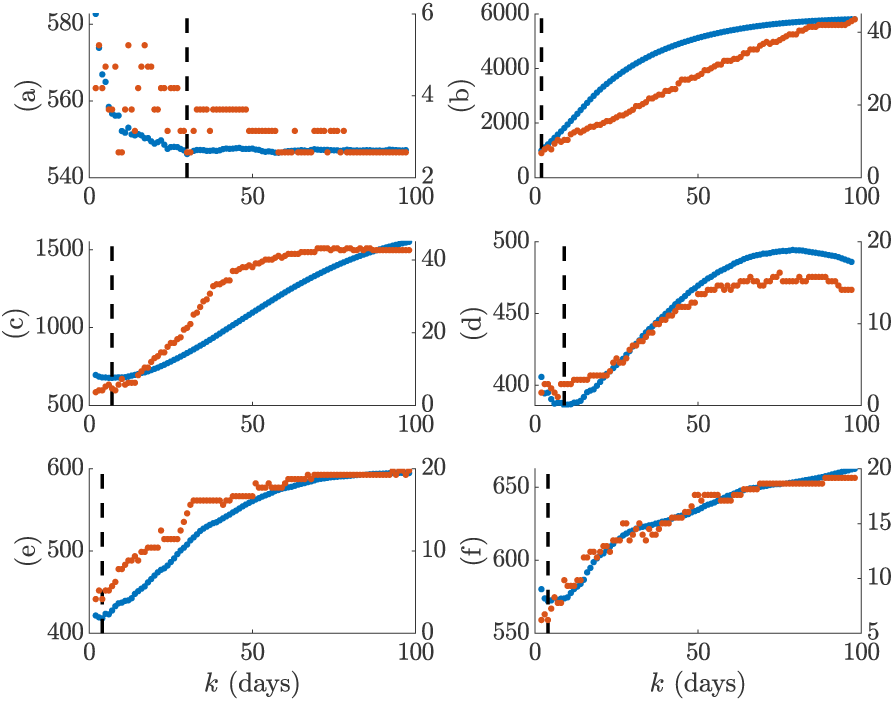
APE prediction accuracy. We compare the APE metric (solid blue, left y axis) to the percentage of true incidence values, *I*_*s*+1_ that fall outside the 95% prediction intervals of 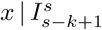 (dotted red, right y axis) over the window search space *k*. The dashed line is *k*\* and panels correspond to those of Fig. 2.

### Diagnosing the Efficacy of Control

A central goal of real-time epidemic forecasting is the rapid and reliable diagnosis or verification of the efficacy of implemented control measures [4]. Knowing whether an intervention is working or not can inform preparedness and resource allocation. Estimating whether *R*_*s*_ < 1 or *R*_*s*_ ≥ 1 is one simple means of assessing efficacy. Effective control measures lead to the former. Here we investigate how the APE metric responds to swift swings in reproduction number and quantify our results over 10^3^ simulated epi-curves with *t* = 150 days. We focus on step-changes in *R*_*s*_ and examine how the successively optimal window length, *k*\*(*s*), computed with data up to time *s*, responds. If *k*\*(*s*) is sensitive to these changes then it is likely a dependable means of diagnosing control efficacy. Note that until now, the optimal window length was *k*\* = *k*\*(*t*).

We consider four possible models: (a) an epidemic that is increasing and uncontrolled, (b) an outbreak that is rapidly controlled, (c) an epidemic with ineffective control that features a late-stage increase in transmission and (d) a two-step control scheme in which the first step was only partially effective. Our results are in Fig. 4 with top graphs overlaying true and predicted incidence across time and middle ones showing the best 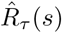 mean estimates under the final *k*\*(*t*). Here we see that the APE-selected model is able to properly distinguish significant changes from stable periods in all scenarios. Model (a) is the most difficult to infer as the increasing reproduction number does not visibly change the shape of the incidence curve.

**Figure 4:**
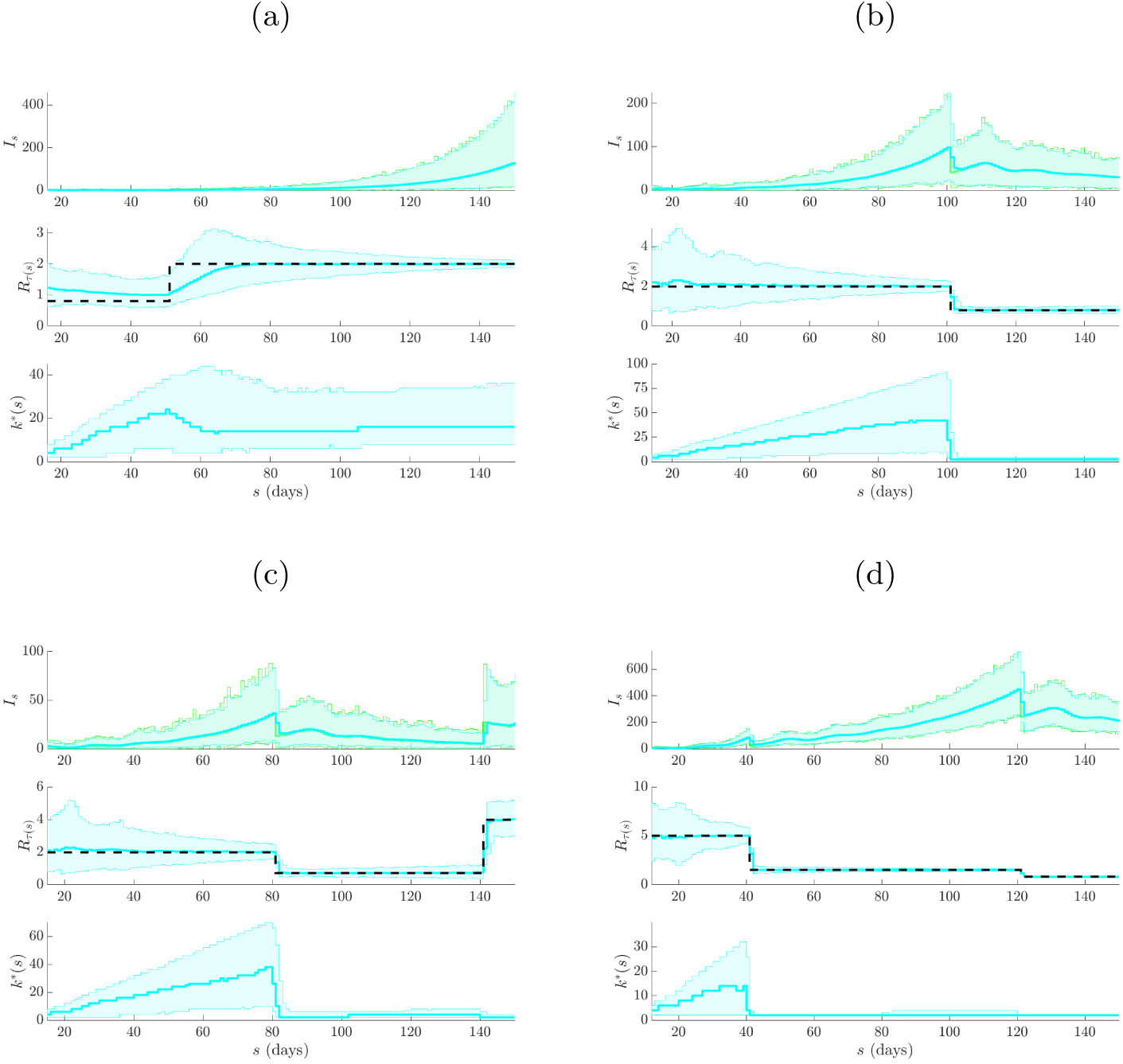
APE sensitivity to real-time transmission changes. We simulate 10^3^ independent incidence curves under renewal models with reproduction numbers indicating (a) increasing, (b) controlled, (c) recovering and (d) cumulatively controlled epidemics. In each panel the top graphs give the true (green) and predicted (cyan) incidence ranges curves, the middle ones provide the estimate of *R*_*s*_ under the final *k*\*(*t*) and the bottom graphs illustrate how the successive *k*\*(*s*) choices of the APE metric vary across time. We find that *k*\*(*s*) responds rapidly to significant changes in the shape of the incidence curves. This makes it sensitive to important fluctuations in *R*_*s*_, especially those that force *R*_*s*_ < 1. The APE metric is suitable for real-time applications and particularly useful for diagnosing the efficacy of control measures.

The bottom graphs of Fig. 4 show *k*\*(*s*) sequentially over time. Here we observe how the APE metric uses data to justify its choices. As data accumulate under a stable reproduction number APE increases *k*\*. This makes sense as there is increasing support for stationary epidemic behaviour. This is especially obvious in Fig. 4b. However, on facing a step-change the APE immediately responds by drastically reducing *k*\*(*s*) – both in scenarios where *R*_*s*_ is changing from above to below 1 and vice versa. This reduction is less visible in Fig. 4a because the observed incidence curve is not as dramatically altered as in Fig. 4b– Fig. 4d. This rapid response recommends the APE as a suitable and sensitive diagnostic of control efficacy.

We illustrate the impact of change-times on the actual APE values at different window lengths in Fig. 5. There we find that when the epi-curve is reasonably stable there is not much difference between the performance at various window lengths and so the APE curves are neighbouring. However, when a non-stationary change occurs there is a clear unravelling of the curves and potentially large gains to be made by optimising *k*. Failing to properly select *k* here can potentially mislead the characterisation of intervention efficacy. This increases the impetus for formal *k*-selection metrics such as the APE.

**Figure 5:**
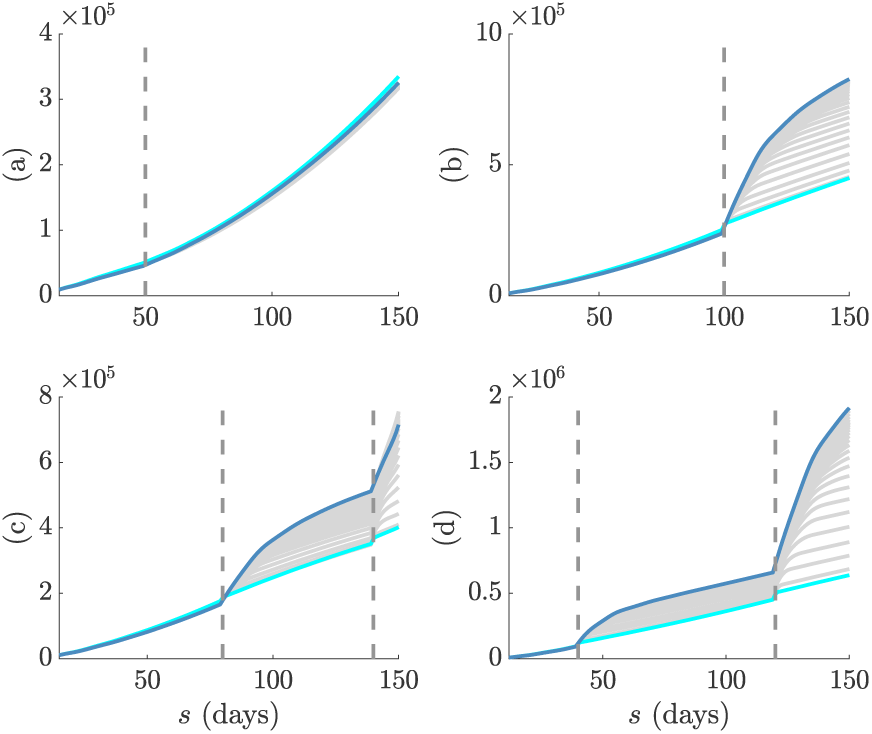
APE predictive scores with time. The successive (in time) APE scores for each window length, *k*, are shown in grey. The scores for the smallest and largest *k* are in cyan and magenta respectively. Graphs correspond to models (a)-(d) in Fig. 4. Dashed lines are the change-times of each model. In (a) the change-time does not significantly affect the epi-curve shape and so the APE scores are close together. In (b)-(d) the change-times notably alter the epi-curve shape so that window length choice becomes critical to performance.

### Empirical Examples

We test our APE approach on two well-studied empirical epidemic datasets: (a) pandemic H1N1 influenza in Baltimore from 1918 [17] and (b) SARS in Hong Kong from 2003 [18]. For each epidemic we extract the incidence curve, total infectiousness and generation time distributions from the EpiEstim R package [8] [19] and apply the APE metric over 2 ≤ *k* ≤ *t/*2 with *t* as the last available incidence time point. We compare the one-step-ahead *I*_*s*+1_ prediction fidelity and the *R*_*τ*(*s*)_ estimation accuracy obtained from the renewal model under the APE-selected *k*\* to that from the model used in [8], which recommended weekly windows (i.e. *k* = 7) after visually examining several window lengths. Our main results are given in Fig. 6 and Fig. 7. We benchmarked our estimates against those directly provided by EpiEstim to confirm our implementation and restrict *k* > 1 to avoid model identifiability issues [9] [20].

**Figure 6:**
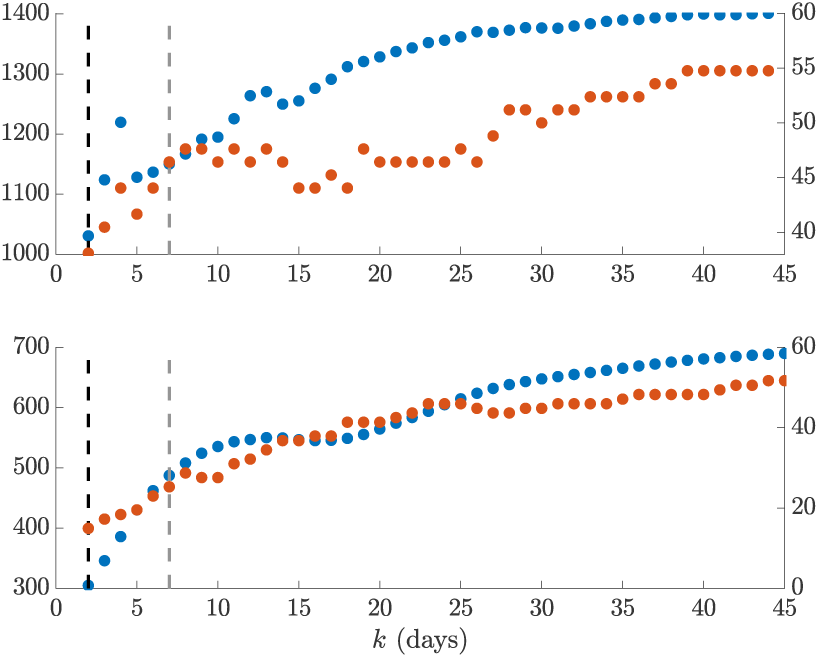
Empirical prediction accuracy. We compare the APE metric (solid blue, left y axis) to the percentage of true incidence values, *I*_*s*+1_, which fall outside the 95% prediction intervals of 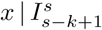 (dotted red, right y axis) across the window search space *k*. The dashed line gives *k*\* (black) and *k* = 7 (grey). The top graph presents results for the influenza 1918 dataset, while the bottom one is for SARS 2003 data. We find that heuristic weekly windows lead to appreciably larger forecasting error than the APE selections.

**Figure 7:**
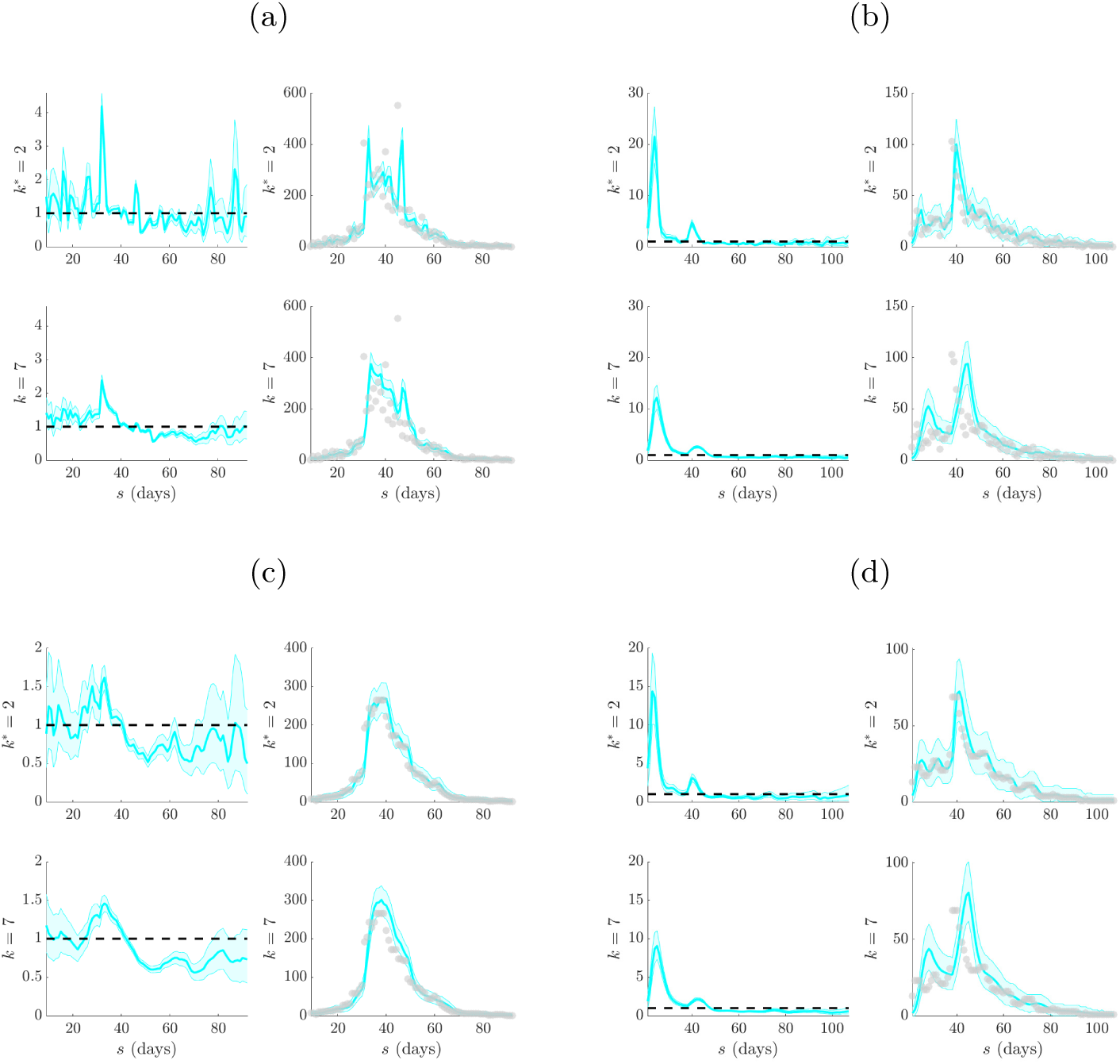
Selection for empirical datasets. Left graphs of each panel compare 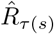 estimates (cyan with 95% confidence intervals) at APE window length *k*\* to those when *k* = 7 (weekly sliding windows). Right graphs give corresponding one-step-ahead predicted incidence curves (cyan with 95% prediction intervals). Dashed lines are the *R* = 1 threshold (left) and dots are the true incidence counts (right). The panels examine epi-data from outbreaks of (a) pandemic influenza (1918) and (b) SARS (2003), which are available in EpiEs-tim [8]. Panels (c) and (d) repeat (a) and (b) respectively, but smooth outliers in the incidence data using 5-day moving averages as in [17]. Interestingly, while *k*\* leads to less believable reproduction number fluctuations, these are justified by the available data in both smoothed and un-smoothed cases. This hints that a Poisson renewal model may not be the most appropriate for these data. The longer weekly windows, which are heuristic recommendations of previous analyses [8], miss important characteristics of the epi-curves.

Intriguingly, we find *k*\* = min *k* = 2 for both scenarios. This yields appreciably improved prediction fidelity (i.e. the coverage of observed incidence values by the 95% prediction intervals), relative to the weekly window choice, as seen in Fig. 6. Between 8 − 10% of incidence data points are better covered by using *k*\* over *k* = 7. This improvement is apparent in the right graphs of (a) and in Fig. 7, where the *k* = 7 case produces stiffer incidence predictions that cannot properly reproduce the observed epi-curve. Weekly windows misjudge the SARS epidemic peak, predict a multimodal SARS incidence curve that is not reflected by the actual data and systematically underestimate influenza case counts when it matters most (i.e. around the high incidence phase).

However, the smaller *k*\* window results in noisier *R*_*τ*(*s*)_ estimates (left graphs of (a) and (b) in Fig. 7), which may be questionable. These rapidly fluctuating reproduction numbers likely motivated the adoption of weekly windows in previous analyses. While the APE-based estimates are indeed more uncertain (a consequence of shorter windows), we argue that they are formally justified by the available data, especially given the weaker predictive capacity of weekly windows. While the noisier *R*_*τ*(*s*)_ estimates do not mislead our understanding of the efficacy of implemented control measures (it is still clear that the influenza epidemic is only partially under control between 40 ≤ *s* ≤ 65 days before recovering, while the SARS outbreak is largely prevented from increasing from *s* > 50 days), they likely overestimate the peak transmissibility of these diseases.

We might suspect that certain artefacts of the data could resolve this issue, rendering a more believable combination of estimated reproduction number and predicted incidence. Particularly, the influenza data seems considerably more affected by the smaller look-back windows. These *k*\* = 2 windows are needed to help predictions get close to the peak incidence values of the data. However, these peaks seem reminiscent of outliers and in the original analysis of [17] they were attributed to possible recollection bias in patients that were questioned. Removing these biases might be expected to lead to smoother APE-justified reproduction numbers.

However, in (c) and (d) of Fig. 7 we find that applying simple 5-day moving averages, as in [17], to ameliorate outliers still does not support a *k*\* > 2 in either scenario, though smoother estimates do result. Weekly windows are still too inflexible to properly predict this averaged incidence. While other signal processing techniques, such as using autocorrelated smoothing prior distributions instead of independent gamma ones over the reproduction numbers, could be applied to test if *k*\* > 2 is justifiable we consider this beyond the scope of this work and somewhat biologically unmotivated. Instead we conjecture that these results support the interesting alternative hypothesis that the epi-curve data are not Poisson distributed.

It is likely that influenza and SARS epi-curves could be better modelled with a less restrictive renewal model. The Poisson renewal model forces a mean-variance equality that may limit the ability to properly predict incidence data sequentially. This limit would then have to be circumvented by APE through the use of smaller sized windows. Using a negative binomial renewal model, which implements the relationship 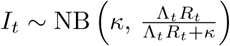, with *κ* as a noise parameter, would relax this equality. The development of APE-based metrics for these generalised models will form a subject of upcoming work. The *κ* parameter should better accommodate transmission heterogeneities and super-spreading events, which are known to affect both influenza and SARS datasets [18] [6].

## Discussion

Inferring the dynamics of the effective reproduction number, *R*_*t*_, in real-time is crucial for forecasting transmissibility and the efficacy of implemented control actions [4]. Renewal models provide a popular platform for prospectively estimating these reproduction numbers, which can then be used to generate incidence projections [16] [6] [3]. However, the dependence of these estimates and predictions on the look-back window size, *k*, which determines the renewal model dimensionality, has never been formally investigated. Previous methods for selecting *k* have generally been heuristic or ad-hoc [8]. Here we have defined and validated a new, rigorous, information-theoretic approach for optimising *k* to available epidemic data.

Our approach was founded on deriving an analytical expression for the renewal model posterior predictive distribution. Integrating this into the general APE metric of [10] led to Eq. (5), a simple, easily-computed yet theoretically justified method for *k*-selection that balances estimation fidelity with prediction accuracy. This APE approach not only accounts for parametric complexity and is formally linked to MDL and BMS [14], but it also has several desirable properties that render it suitable for handling the eccentricities of real-time infectious disease applications, as follows:

a. Small outbreak sample size. Early-on in an epidemic data are scarce and uncertainty is large. The APE approach is valid and computable at all sample sizes at which the renewal model is identifiable and hence is applicable to emerging outbreaks [11]. Its emphasis on using the full predictive distribution and its Bayesian formulation allow it to properly account for large uncertainties and to handle different (prior) hypotheses about *R*_*t*_, as the epidemic progresses from small to large data regimes.
b. Non-stationary transmission. Incidence time series are not independent and identically (iid) distributed [6]. Instead they are sequential, autocorrelated and possess non-stationary (time-varying) statistics. The APE formally accounts for these properties. Leave-one-out cross validation (CV) is a popular model selection approach that is related to APE. It can be defined as 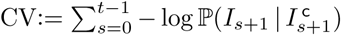, where 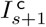 means that all incidence points except *I*_*s*+1_ are included [21]. Comparing this to Eq. (9) (see Methods) we see that APE is a modified CV that is explicitly specialised for non-stationary, accumulating time-series.
c. Lack of a ‘true model’. It is highly unlikely that the underlying *R*_*t*_ is truly described by one of the piecewise-constant functions assumed within renewal models. Unlike many model selection criteria, the APE does not require a true model to exist within the set being evaluated [14]. It is only interested in finding the model that best predicts the data and so emphasises accurate forecasting and data-justified complexity. However, should a true model exist, the APE is statistically consistent i.e. as data accumulate it will select the data-generating model with probability 1 [14].

We tested our metric on both simulated and empirical datasets. We started by examining various distinct, time-varying reproduction number profiles in Fig. 2, for which no true model existed. By comparing the APE-optimised *k*\* against long and short window sizes, we found that not only does APE meaningfully balance *R*_*t*_ estimates to achieve good prediction capacity, but also that getting *k* wrong could promote strikingly different conclusions from the same dataset. This behaviour held consistent for outbreaks with both small and large numbers of infecteds.

We then investigated several step-changing reproduction number examples in Fig. 4 to better expose the underlying mechanics of our approach, and to showcase its performance on epidemics with strong non-stationary transmission. These examples featured rapid changes, also known as events in information theory, caused by effective and ineffective countermeasures [4]. In all cases, our metric responded rapidly whilst maintaining reliable incidence predictions in real-time. Such rapid, event-triggered responses (as in Fig. 5) are known to be time-efficient [22]. A key objective of short-term forecasting is the speedy diagnosis of intervention efficacy. Fig. 4 confirmed the practical value of our method towards this objective.

While our metric is promising, we caution that much work remains to be done. Standard Poisson renewal models, such as those we have considered here, make several limiting assumptions including that (i) all cases are detected (i.e. there is no significant sampling bias), (ii) the serial interval and generation time distribution coincide and do not change over the epidemic lifetime and (iii) that heterogeneities in transmission within the infected population have negligible effect [8]. Our analysis of H1N1 influenza (1918) and SARS (2003) flagged some of these concerns. Comparing the *k*\* to previously recommended weekly windows led to some interesting revelations.

The APE approach selected a notably shorter window (*k*\* = 2 days) for both datasets. While this produced noisy *R*_*s*_ estimates that seemed less reliable than those obtained with weekly windows, the predicted incidence values were central to understanding this discrepancy. The weekly windows were considerably worse at forecasting the observed epi-curve, often systematically biased around the peak of the epidemic and sometimes predicting multi-modal curves that were not reflected by the existing data. From a model selection perspective, the shorter windows are therefore justified.

However, the noisy *R*_*t*_ estimates are still undesirable. Both datasets are known to potentially contain super-spreading heterogeneities and other biases that may lead to outliers [17] [8]. Moving-average filters were applied in [17] to remove these artefacts from the H1N1 data. We used the same technique to regularise both datasets and then re-applied the APE. Results remained consistent, providing evidence for the use of more flexible negative binomial renewal models, which can better handle these biases. In an upcoming study we will adapt the APE for these more general renewal methods.

Real-time, model-based epidemic forecasts are quickly becoming an integral prognostic for resource allocation and strategic intervention planning [23]. As global infectious disease threats elevate, model-supported predictions, which can inform decision making, have become imperative. However, much is still unknown about the fundamentals of epidemic prediction and this uncertainty has inspired some reluctance in public health applications [23]. As a result, recent studies have called for careful evaluation of the predictive capacity of models, and demonstrated the importance of having standard metrics to compare models and qualify their uncertainties [24] [25].

Here we have presented one such metric. Short-term forecasts aid immediate decision making and are more actionable than long-term ones (reliability decreases sharply with prediction horizon) [24] [23]. While we focussed on renewal models, the general APE metric of Eq. (9) can select among diverse types of models, facilitating comparative evaluation of their short-term predictive capacity and reliability. Its direct use of posterior predictive distributions within an honest, causal framework allows proper and problem-specific inclusion of uncertainty [11]. Common metrics such as the mean absolute error lack this adaptation [24]. Given its information-theoretic links and demonstrated performance we hope that our APE-based method can serve as a rigorous benchmark for model-based forecasts.

## Methods

### Epidemic Renewal Models

Consider an incidence curve over times 1 ≤ *s* ≤ *t* for an epidemic with effective reproduction number and total infectiousness at *t* of *R*_*t*_ and Λ_*t*_. While *R*_*t*_ describes the branching of the epidemic (the number of secondary cases originating from a primary one), 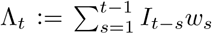, controls how infected cases propagate, via the generation time distribution, which is defined by *w*_*s*_. Here *w*_*s*_ is the probability that a primary case takes between *s* − 1 and *s* days to generate a secondary case [5]. This distribution is intrinsic to a disease and 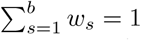 for some memory time *b*. We make the common assumptions that the generation time distribution is known and does not change with time [8].

The quantities *R*_*t*_ and Λ_*t*_ therefore completely describe the transmissibility of an epidemic, an idea formalised by the renewal model [5]. Generally *R*_*t*_ is unknown and must be inferred from 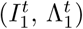. Its log-likelihood under the standard renewal model, 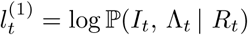 follows as in Eq. (6) [6].

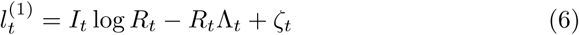

Here *ζ*_*t*_ = − log *I*_*t*_!+*I*_*t*_ log Λ_*t*_ collects terms that do not depend on the parameter *R*_*t*_. The superscript of 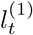 highlights that this model employs a unit window length, and hence only uses (*I*_*t*_, Λ_*t*_) to infer *R*_*t*_. While this construction maximises model flexibility, the resulting estimates can be noisy and over-fitting is likely [6] [9]. Grouping is therefore employed [8].

This assumes that the reproduction number, denoted *R*_*τ*(*t*)_, is constant over the past *k* time units [8], and leads to a piecewise-constant function that classifies between meaningful and negligible reproduction number changes. The grouped log-likelihood, 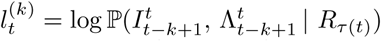, with parameter-independent term *ζ*_*τ*(*t*)_ = ∑_*s*∈*τ* (*t*)_ − log *I*_*s*_! + *I*_*s*_ log Λ_*s*_ and grouped sums *i*_*τ*(*t*)_ : = ∑_*s*∈*τ* (*t*)_ *I*_*s*_ and *λ*_*τ*(*t*)_ := ∑_*s*∈*τ* (*t*)_ Λ_*s*_ follows in Eq. (7). At *k* = 1 we recover Eq. (6).

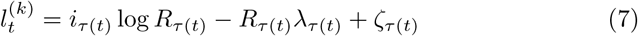

The maximum likelihood estimates (MLEs) and Fisher information (FI) of Eq. (7) provide insight into the benefits of *k*-grouping. The MLE facilitates unbiased inference, while the FI bounds our confidence around the MLE (it measures the inverse of estimate variance) [27]. The MLE, 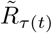, is the solution to 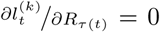, while the FI is 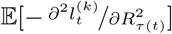 [27]. We actually compute the FI for the square root of 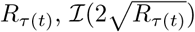, as it is known to have optimal properties [20]. Eq. (8) then follows [9].

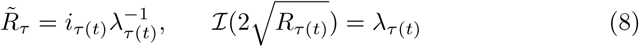

Comparing Eq. (8) to equivalent expressions when *k* = 1 reveals the impact of grouping. This gives 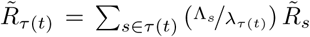 and 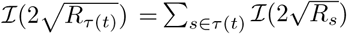. The grouped MLE is hence a weighted moving average of the ungrouped MLEs, which explains why noise is reduced. The grouped FI is a linear summation of the ungrouped FIs, implying that estimate precision also increases with grouping. Unfortunately, these advantages come at the expense of elevated tracking bias. At the extreme of *k* = *t*, for example, 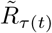 is a stable *t*-point average, that can only be gradually perturbed by new incidence data. Thus, we trade the sensitivity to rapid *R*_*s*_ changes for smaller estimate variances [8]. The need to formally mediate this trade motivated the APE metric.

### Prospective Model Selection

In [20] a minimum description length (MDL) solution was proposed for retrospectively selecting a different but related *k* that defines the non-overlapping window size optimising historical reproduction number estimates. This method, by exploiting an often neglected aspect of model complexity, known as parametric complexity [14], was able to outperform standard measures such as Akaike (AIC) and Bayesian information criteria (BIC). While this MDL technique is not directly applicable here, as prospective performance requires a different optimisation [15], we heed the lesson about accounting for parametric complexity.

The criteria we propose is the APE [10], which values models on their ability to generalise i.e. predict unseen data from the generating process [11]. Practically, this is implemented by sequentially predicting the data observed at time *s* + 1 (i.e. one-step-ahead of *s*) using the subset of data preceding it [15]. This means that we causally predict *I*_*s*+1_ at every *s* given a *k*-window back in time of *τ* (*s*) = {*s, s* − 1, …, *s* − *k* + 1}. This window is truncated if *s* < *k* so that min *τ* (*s*) ≥ 1. We then evaluate our prediction (e.g. the posterior mean *Î*_*s*+1_) against the observed *I*_*s*+1_. The *k* minimising the cumulative one-step-ahead prediction error up to the present *t*, which we term *k*\*, gives the renewal model that best predicts the unseen datum at *t* + 1.

We generally do not use *Î*_*s*+1_ directly, but instead obtain its full predictive distribution 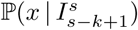, with *x* as some value of the predicted incidence at time *s* + 1. Eq. (9) then defines the APE as a cumulative log-score function.

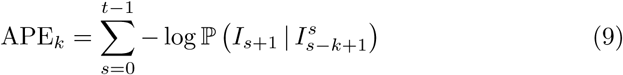

The optimal window, *k*\* := arg min_*k*_ APE_*k*_, is easy to compute provided the predictive distribution in Eq. (9) is calculable. Fig. 1 summarises and illustrates the APE approach. By using the full posterior predictive distribution the APE appropriately accounts for predictive uncertainty and is specialised to the problem of interest. A point-estimate alternative to APE, known as predictive mean squared error (PMSE), can be used when this distribution is not available [10] and is defined as 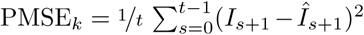. While the PMSE is not as tailored to specific problems, both metrics converge when errors are normally distributed [11]. Other score functions can be used with these one-step-ahead predictions when application-specific insights are available [28].

The APE metric has formal links to MDL and Bayesian model selection (BMS). BMS also includes parametric complexity and is asymptotically equivalent to the MDL when Jeffreys prior is used within the BMS [13] [14]. Interestingly, because any joint distribution can be decomposed as 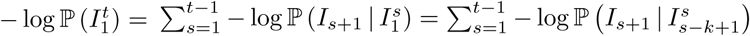, APE, under certain regularity conditions, is equivalent to both of these measures [29]. The latter equality is from the finite memory of the renewal model, which depicts the non-stationary nature of epidemics.

However, the APE is simpler and more transparent, requiring no difficult integral evaluations [9]. Consequently, the APE not only accounts for parametric complexity (implicitly), but also applies to models of arbitrary complexity [11]. The drawbacks of APE are that it requires the data to be ordered in time, and being data-driven, its computational complexity increases linearly in both the number of models to be assessed and the size of *t* [30]. Overall, APE provides a simple and optimal solution to window selection, which surprisingly has not penetrated the epidemiological literature.

## Acknowledgments

KVP and CAD acknowledge joint Centre funding from the UK Medical Research Council and Department for International Development under grant reference MR/R015600/1. CAD thanks the UK National Institute for Health Research Health Protection Research Unit (NIHR HPRU) in Modelling Methodology at Imperial College London in partnership with Public Health England (PHE) for funding (grant HPRU-2012–10080).

## Author Contributions

KVP designed and performed research and wrote the initial draft of the paper. CAD supervised the research. All authors edited and approved the final draft.

